# Genome sequencing of the Monkeypox virus 2022 outbreak with Amplicon-based Oxford Nanopore MinION sequencing

**DOI:** 10.1101/2022.10.20.512862

**Authors:** Annika Brinkmann, Katharina Pape, Steven Uddin, Niklas Woelk, Sophie Förster, Heiko Jessen, Janine Michel, Claudia Kohl, Lars Schaade, Andreas Nitsche

**Author notes:** Address correspondence to Annika Brinkmann.

## Abstract

We present an amplicon-based assay for MinION Nanopore sequencing of monkeypox virus genomes from clinical specimens, obtaining high-quality results for up to 99 % of the genomes, even for samples with high Ct values of up to 30 which are not suitable for shotgun sequencing.

## Introduction

With the first confirmed cases of the 2022 multi-country outbreak of monkeypox virus (MPXV), laboratories all over the world have started virus genome sequencing. The first genome was presented by a laboratory in Portugal on 28 May 2022, followed by many countries all over the world (https://virological.org/). As of 11 October 2022, 2319 genomes have been sequenced from 71,096 known cases (3.3 %). In contrast to sequencing efforts at the beginning of and during the current SARS-CoV-2 pandemic, the number of MPXV genomes being uploaded to public databases is quite low. Sometimes there are no new genomes being added on several consecutive days, and few laboratories participate regularly in sequencing (https://nextstrain.org/monkeypox/hmpxv1?label=clade:B.1).

MPXV, a species of the family *Poxviridae* and genus Orthopox virus, which includes the eradicated Variola virus, has been known to be endemic in central (clade I) and western (clade IIa) African countries. MPXV of the 2022 outbreak has been assigned recently as clade IIb by the WHO, being phylogenetically closest related to MPXV cases which have been imported from Nigeria to non-endemic countries such as the United Kingdom, Israel and Singapore, in 2018, 2019 and 2021 (1-4). However, in contrast to previous outbreaks, the comparably high number of genomic sequences has revealed an unexpected high number of mutations in the MPXV genome of the current outbreak, and the mode and length of human transmission chains seem to have changed (5).

Sequencing is inevitable to explain these occurrences in the current outbreak. The requirement of technical laboratory skills and necessities (e.g. availability of biosafety level 3 laboratories for virus propagation) and access to sequencing platforms with high sequencing output, such as Illumina, might impede frequent sequencing for many laboratories. Furthermore, the size (197,000 bp) and complexity of the MPXV genome with many repeat regions and possible deletions and insertions require poxvirus-specific bioinformatics expertise and make completely automated analysis, as is often used for SARS-CoV-2, difficult.

Here, we present an amplicon-based assay for MinION nanopore sequencing, which can scale up sequencing for laboratories already performing shotgun sequencing and can enable sequencing for laboratories without access to Illumina platforms. We could show that up to 99 % of the MPXV genomes could be generated with sequencing of only 200,000 reads, also those of samples with higher Ct values of up to 30 which are challenging to sequence on Illumina platforms and require an extensive number of reads. We could prove the accuracy of the approach by comparing the generated genomes with high-quality Illumina-derived genomes from the same samples. We propose that amplicon-based Nanopore sequencing can contribute to solve unanswered questions of the 2022 outbreak and monitor the frequency and appearance of mutations as the outbreak continues.

## Methods

### Primer Design and Evaluation

Using a tilted approach for primer design, 682 primer pairs for 375 bp amplicons were designed for the amplification of MPXV, using Monkeypox/PT0006/2022|sampling_date_20220515 (accession ON585033.1) (https://ampliseq.com/browse.action).

### PCR amplification, sequencing and analysis

The patient samples were amplified in a single reaction with the following PCR conditions: 3 µl of viral DNA, 5 µl of primer pool, 2 mM dNTP (Invitrogen, Karlsruhe, Germany), 2 µl of 10 × Platinum Taq buffer, 4 mM MgCl_2_ and 2 µl of Platinum Taq polymerase (Invitrogen) with added water to a final volume of 25 µl. Cycling conditions were 94°C for 2 min followed by 35 amplification cycles at 94°C for 20 s and 60°C for 4 min.

The amount of human DNA in relation to MPXV DNA in each sample was determined with a generic Orthopox virus qPCR (rpo) and a PCR that detects human nucleic acid (*MYC*) (6, 7). Amplified samples were processed for nanopore sequencing on the MinION (Oxford Nanopore Technologies, Oxford, United Kingdom) with the ligation sequencing kit 1D, SQK-LSK109 (Oxford Nanopore Technologies). Samples were sequenced on different flow cells with at least 200,000 reads per sample. The bioinformatics analysis was performed as described elsewhere (8).

## Results

### Comparison of MPXV genomes sequenced by Illumina and amplicon-based Nanopore sequencing

We compared five MPXV genomes sequenced with Illumina platform with the amplicon-based Nanopore sequencing. Figure 1 shows the number and position of mutations (in relation to ON676708 MPXV_USA_2021_MD) and non-covered regions of genomes. All mutations in relation to MPXV ON676708 were called concordant for both methods of the corresponding genome pair, with no mutation missed or added. The overall genome coverage of the Nanopore-sequenced samples was slightly lower than the Illumina-sequenced ones, but not lower than 99.69 %. Non-covered regions of all Nanopore-sequenced samples were found at both ends (60 and 180 bp), at 19,156 bp (146 bp) and at 133,056 bp (78 bp).

**FIGURE 1:**
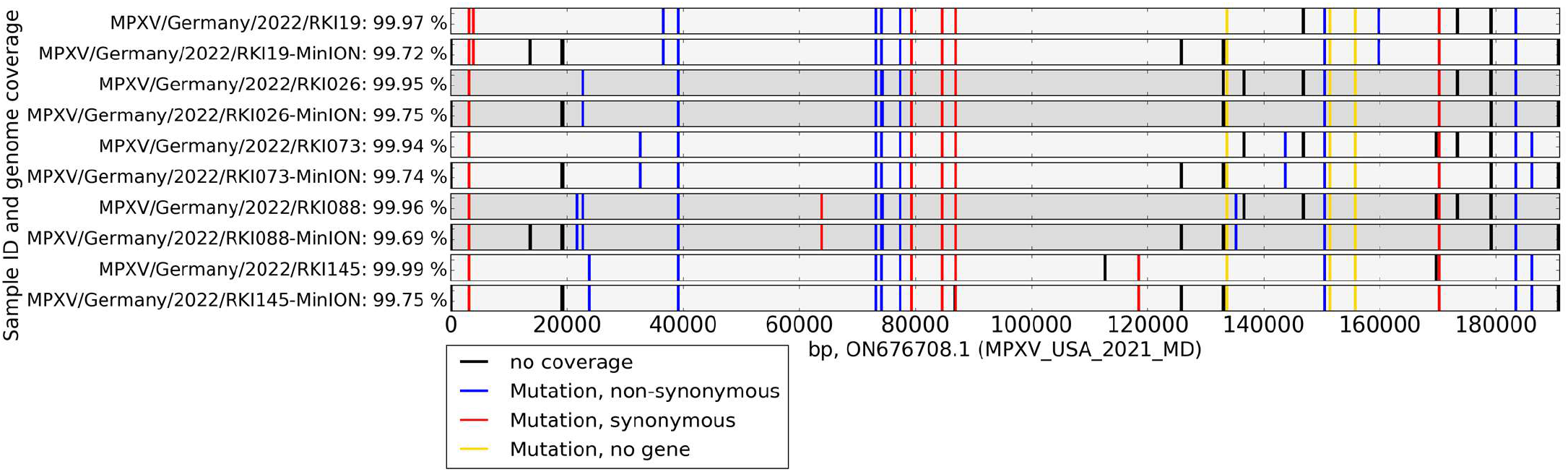
The genomes of MPXV samples sequenced by the Nanopore Amplicon approach (NCBI MPXV/Germany/2022/RKI19-MinION, MPXV/Germany/2022/RKI026-MinION, MPXV/Germany/2022/RKI073-MinION, MPXV/Germany/2022/RKI088-MinION and MPXV/Germany/2022/RKI145-MinION) and Illumina platform (NCBI MPXV/Germany/2022/RKI19, MPXV/Germany/2022/RKI026, MPXV/Germany/2022/RKI073, MPXV/Germany/2022/RKI088 and MPXV/Germany/2022/RKI145) were compared, including mutations and non-covered regions. Missing coverage: black bars; non-synonymous mutations: blue bars; synonymous mutations within genes: red bars; mutation outside of genes: yellow bars.

**FIGURE 2:**
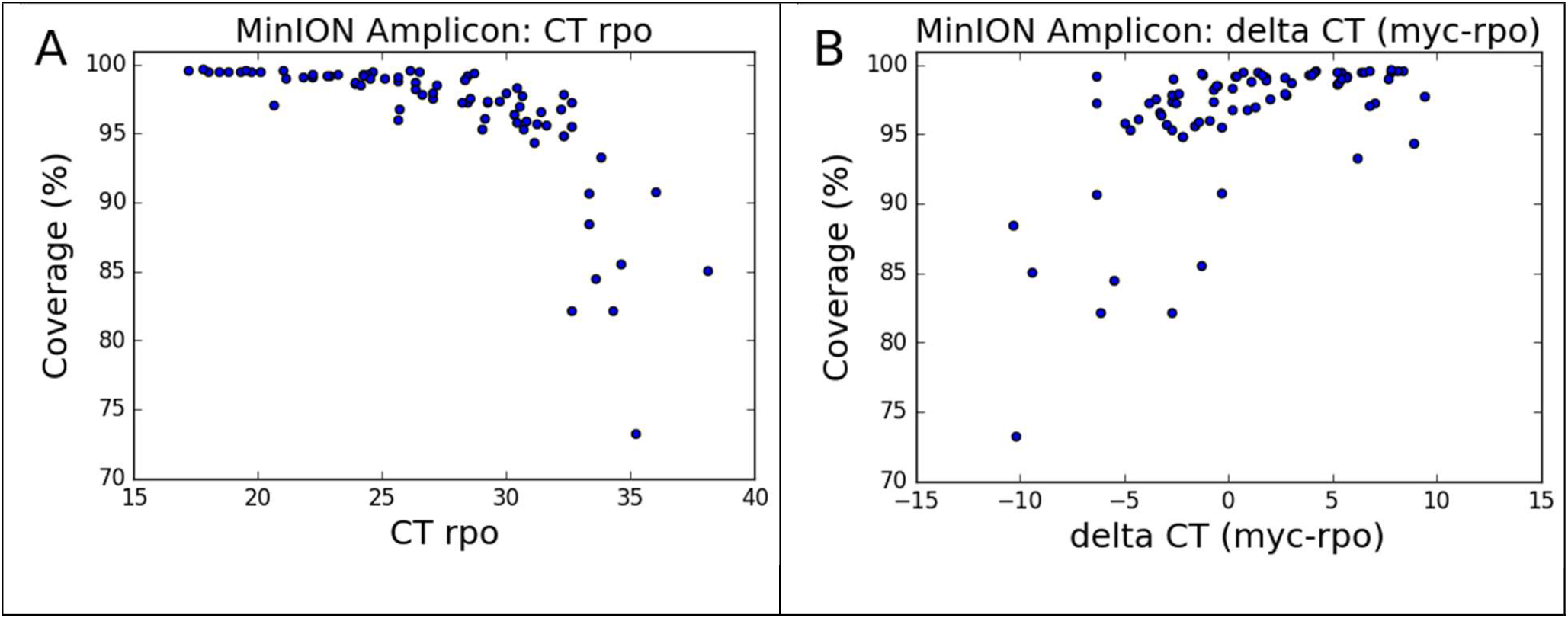
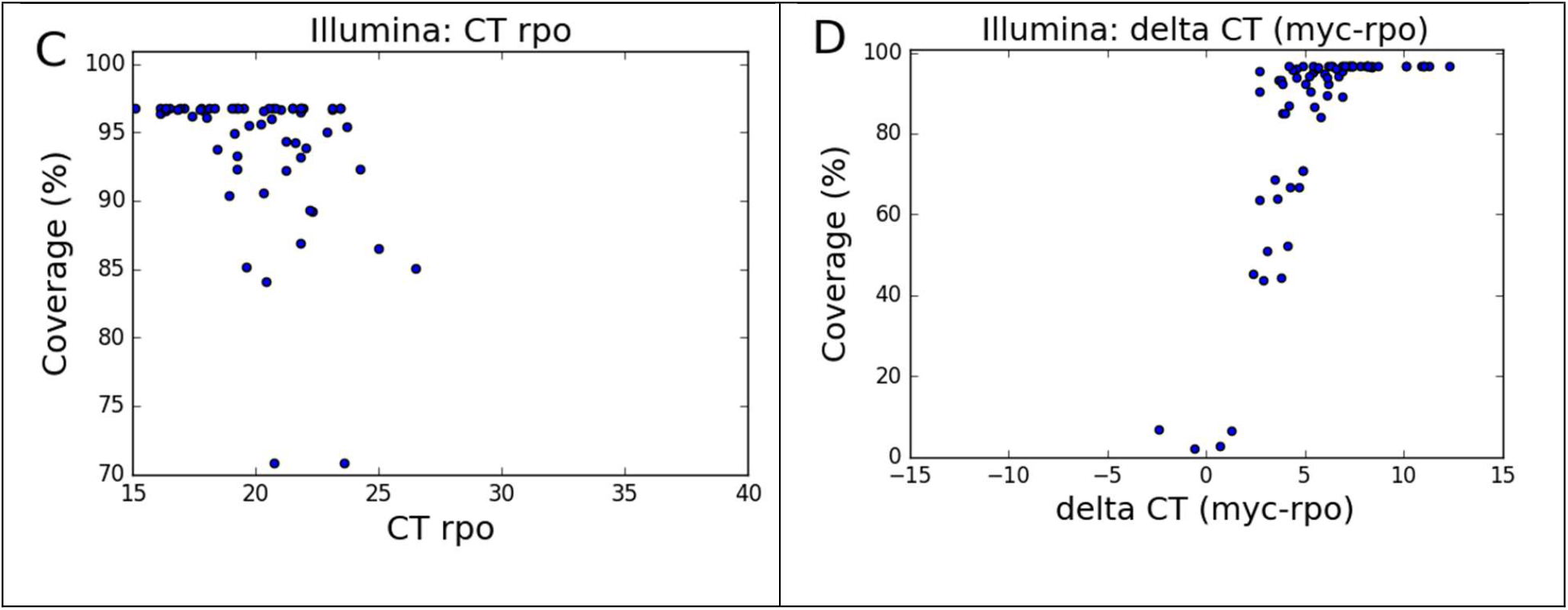
77 patient samples were screened for MPXV by qPCR and sequenced with the Nanopore Amplicon approach. A mean coverage of 96.41 % could be reached for all samples with 200,000 sequenced reads. Up to Ct 25, a mean coverage of 99.19 % could be reached. Unlike Illumina sequencing (shown for a different batch of 77 samples with Cts of up to 27 and shown for 10,000,000 reads), where the resulting genome coverage is mostly correlated to the ratio of human (background) and MPXV DNA in each sample, the genome coverage after amplicon sequencing is merely dependent on the MPXV concentration.

### Screening of patient samples

We screened 77 MPXV patient samples with Cts ranging from 17 to 38 with the amplicon-based Nanopore assay for MPXV. The genome coverage for the sequenced genomes was in correlation with the Ct value (rpo) of the sample. Almost complete coverage from 99.49 to 99.67 % (mean: 99.54) could be reached for Ct values of up to 20, mean coverage of 99.03 % for Ct values from 20 to 25 and mean coverage of 97.9 % for Ct values from 25 to 30. Above 30, a mean coverage of 92.12 % could be reached, with coverage from 98.34 to the lowest coverage of 73.21 for a Ct value of 35.2. For each sample 200,000 reads were evaluated, with higher coverage including all sequenced reads.

## Discussion

Multiple unsolved questions regarding the 2022 MPXV outbreak have been raised, including the mode of transmission, the unexpected high number of mutations in clade IIb and possible ongoing adaption to the human host. Only by sequencing samples from the current outbreak, including multiple samples from the same host, these questions can be resolved (9). However, sequencing is often impeded by a poor ratio of MPXV to human DNA in samples. For shotgun sequencing, even for samples with a relatively high concentration of the virus, the concentration of viral reads in the sample is rarely higher than 0.5 %, making sequencing output of several million reads necessary.

Here we show that amplicon-based sequencing can be used to upscale sequencing, while only 200,000 reads are necessary for full coverage of the genome. The assay can be used for samples with a Ct of up to 25 with mean coverage of 99.19 % and up to Ct 30 with mean coverage of 97.9. Overall coverage of 99.75 % could be reached, with the unavoidable lack of coverage of regions at both ends and short stretches with high GC content and low Tm.

However, one must exercise high caution when reconstructing genomes from amplicon-based sequencing with Nanopore, as uploading genomes with unconfirmed mutations should be avoided. It is necessary to mark the consensus sequences at regions with coverage at least below depth < 10 and to confirm that no called mutations are the outcome of sequencing errors. Furthermore, MPXV genomes can have large insertions and deletions which in most cases will be missed by amplicon-based sequencing (10). The MPXV core genome is conserved, whereas the termini contain genes related to host adaption. In Orthopox viruses the deletion of these genes has been shown to be a cause of adaption to the host (11). Significant deletions have been shown in the MPXV genomes from the current outbreak, possibly an adaption to the human host (12). MPXV genomes generated by amplicon sequencing with a coverage below 99 % should therefore be checked for uncommon stretches of N’s, where possible indels could be missed, and then be subjected to high-quality shotgun sequencing.

## Conclusion

We have shown that amplicon-based sequencing can be used to reconstruct high-quality, accurate MPXV genomes. Although shotgun sequencing is necessary in some cases, especially to identify insertions and deletions in the MPXV genome, we propose that amplicon-based sequencing can be used to track single nucleotide polymorphisms and to screen samples for possible cases of insertions and deletions.

## Acknowledgments

We kindly thank Ute Kramer for technical assistance and Ursula Erikli for copy-editing. The authors are grateful to Andrea Thürmer and the MF2 team for Illumina sequencing service.

## Ethics statement

The studies involving human participants were reviewed and approved by the Ärztekammer Berlin (Berlin Medical Association; #Eth-44/22). The patients/participants provided their written informed consent for sequencing.

## Conflict of interest

None declared.

## Authors’ contributions

Annika Brinkmann designed the assay, analyzed the data and wrote the manuscript. Katharina Pape, Steven Uddin, Niklas Woelk and Sophie Förster analyzed the samples and the data. Heiko Jessen provided clinical specimens. Janine Michel and Claudia Kohl analyzed the suspected MPXV specimens. Lars Schaade and Andreas Nitsche conceptualized the approach and wrote the manuscript.

## Notes

### Competing Interest Statement

The authors have declared no competing interest.

